# Cell competition between wild-type and JAK2V617F mutant cells prevents disease relapse after stem cell transplantation in a murine model of myeloproliferative neoplasm

**DOI:** 10.1101/2021.08.24.457589

**Authors:** Haotian Zhang, Melissa Castiglione, Lei Zheng, Huichun Zhan

## Abstract

Disease relapse after allogeneic stem cell transplantation is a major cause of treatment-related morbidity and mortality in patients with myeloproliferative neoplasms (MPNs). The cellular and molecular mechanisms for MPN relapse are not well understood. Here, we established a murine model of MPN relapse, in which ∼60% of the MPN recipient mice develop disease relapse after receiving stem cell transplantation with wild-type marrow donor. Using this model, we find that impaired wild-type cell function is associated with MPN disease relapse. We also show that competition between wild-type and JAK2V617F mutant cells can modulate the immune cell composition and PD-L1 expression induced by the JAK2V617F oncogene. These results suggest that cell competition between wild-type donor cells and JAK2V617F mutant recipient cells can prevent MPN disease relapse after stem cell transplantation.

## INTRODUCTION

Myeloproliferative neoplasms (MPNs) are clonal stem cell disorders characterized by hematopoietic stem/progenitor cell (HSPC) expansion, overproduction of mature blood cells, and an increased risk of transformation to acute leukemia. The acquired kinase mutation JAK2V617F plays a central role in most patients with these disorders. The HSPC compartment in MPN is heterogeneous with the presence of both JAK2 wild-type and JAK2V617F mutant cells in most patients with MPN^1^. Despite mutant cells bearing an *in vitro* proliferative advantage because of the constitutive kinase activity of JAK2V617F, in many patients, there is little or no change in the mutant/wild-type cell ratio over long periods of follow up^2-4^. In other patients, the MPN can evolve to acute leukemia and patients experience high relapse rates following allogeneic stem cell transplantation, the only curative treatment for patients with MPNs^5-8^. Such disease relapse is seen in up to 40% of patients after stem cell transplantation^5,7,9^ and is a leading cause of transplant-related morbidity and mortality in patients with these diseases. Mechanisms for why the MPN disease relapses in some patients while remains in remission in others are not well understood, partly due to the lack of robust model systems.

Endothelial cells are an essential component of the hematopoietic niche^10^. The JAK2V617F mutation can be detected in microvascular ECs isolated from the liver^11^, spleen^12^, and marrow^13^ of many patients with MPNs. The mutation can also be detected in circulating EC progenitors derived from the hematopoietic lineage and, in some reports based on *in vitro* culture assays, in endothelial colony-forming cells from patients with MPNs^12-16^. Previously, using a transgenic mouse expressing JAK2V617F specifically in all hematopoietic cells and vascular ECs, we found that the JAK2V617F mutant HSPCs are relatively protected by the JAK2V617F-bearing vascular niche from an otherwise lethal dose of irradiation. After stem cell transplantation with wild-type marrow donor, MPN disease relapse occurs in ∼60% of these JAK2V617F-positive recipient mice while the others remain in remission during a 3-month follow up^17^. Although the JAK2V617F mutation is present in all ECs in the transgenic mice while vascular niche in MPN patients is likely heterogeneous with the co-existence of both normal and mutant ECs, this model provides us the unique opportunity to study the mechanisms for MPN disease relapse by comparing HSPC functions between the relapse mice and those in remission. Using this model, we found that cell competition between wild-type and JAK2V617F mutant HSPCs after marrow transplantation can prevent MPN disease relapse.

## Materials and Methods

### Experimental mice

JAK2V617F Flip-Flop (FF1) mice^18^ was provided by Radek Skoda (University Hospital, Basal, Switzerland) and *Tie2-Cre* mice^19^ by Mark Ginsberg (University of California, San Diego). FF1 mice were crossed with Tie2-Cre mice to express JAK2V617F specifically in all hematopoietic cells (including HSPCs) and vascular ECs (Tie2^+/-^ FF1^+/-^, or Tie2FF1), so as to model the human diseases in which both the hematopoietic stem cells and ECs harbor the mutation. All mice used were crossed onto a C57BL/6 background and bred in a pathogen-free mouse facility at Stony Brook University. Animal experiments were performed in accordance with the guidelines provided by the Institutional Animal Care and Use Committee.

### Stem cell transplantation assays

Recipient mice were irradiated with two doses of 540 rad 3 hours apart. Donor cells were injected into recipients by standard intravenous tail vein injection using a 27G insulin syringe. For competitive transplantation, 5×10^5^ CD45.2 donor marrow cells from Tie2FF1 mice were injected intravenously together with 5×10^5^ competitor CD45.1 wild-type marrow cells. For noncompetitive transplantation, 1×10^6^ unfractionated donor marrow cells were transplanted into wild-type recipients by intravenous tail vein injection.

### Complete blood counts

Peripheral blood was obtained from the facial vein via submandibular bleeding, collected in an EDTA tube, and analyzed using a HM5 Hematology Analyzer (Zoetis, Parsippany, NJ).

### Marrow and spleen cell isolation

Murine femurs and tibias were first harvested and cleaned thoroughly. Marrow cells were flushed into PBS with 2% fetal bovine serum using a 25G needle and syringe. Remaining bones were crushed with a mortar and pestle followed by enzymatic digestion with DNase I (25U/ml) and Collagenase D (1mg/ml) at 37 °C for 20 min under gentle rocking. Tissue suspensions were thoroughly homogenized by gentle and repeated mixing using 10ml pipette to facilitate dissociation of cellular aggregates. Resulting cell suspensions were then filtered through a 40uM cell strainer.

Murine spleens were collected and placed into a 40uM cell strainer. The plunger end of a 1ml syringe was used to mash the spleen through the cell strainer into a collecting dish. 5ml PBS with 2% FBS was used to rinse the cell strainer and the resulting spleen cell suspension was passed through a 5ml syringe with a 23G needle several times to further eliminate small cell clumps.

### Flow cytometry

All samples were analyzed by flow cytometry using a FACSAriaTM III or a LSR II (BD Biosciences, San Jose, CA, USA). CD45.1 (Clone A20, BD Biosciences), CD45.2 (Clone 104, Biolegend, San Diego, CA), Lineage cocktail (include CD3, B220, Gr1, CD11b, Ter119; Biolegend), cKit (Clone 2B8, Biolegend), Sca1 (Clone D7, Biolegend), CD150 (Clone mShad150, eBioscience, San Diego, CA), CD48 (Clone HM48-1, Biolegend), PD-L1 (Clone 10F.9G2, Biolegend), CD3 (Clone 17A2, Biolegend), CD4 (Clone GK1.5, Biolegend), CD8 (Clone 53-6.7, Biolegend), B220 (Clone RA3-6B2, Biolegend), CD11b (Clone M1/70, Biolegend), Ly-6C (Clone HK1.4, Biolegend), and Ly-6G (Clone 1A8, Biolegend) antibodies were used.

### BrdU incorporation analysis

Mice were injected intraperitoneally with a single dose of 5-bromo-2′-deoxyuridine (BrdU; 100 mg/kg body weight) and maintained on 1mg BrdU/ml drinking water for two days. Mice were then euthanized and marrow cells isolated as described above. For analysis of HSPC (Lin^-^cKit^+^Sca1^+^) proliferation, marrow cells were first stained with fluorescent antibodies specific for cell surface HSPC markers, followed by fixation and permeabilization using the Cytofix/Cytoperm kit (BD Biosciences), DNase digestion (Sigma, St. Louis, MO), and anti-BrdU antibody (Biolegend) staining to analyze BrdU incorporation^20^. Data were acquired using a LSR II flow cytometer.

### Analysis of apoptosis by active caspase-3 staining

Marrow cells were stained with fluorescent antibodies specific for cell surface HSPC markers, followed by fixation and permeabilization using the Cytofix/Cytoperm kit (BD Biosciences). Cells were then stained using a rabbit anti-activated caspase-3 antibody^20^. Data were acquired using a LSR II flow cytometer.

### Analysis of senescence by senescence associated β-galactosidase (SA-β-Gal) activity

Marrow cells were stained with fluorescent antibodies specific for cell surface HSPC markers. Cells were then washed and fixed using 2% paraformaldehyde and incubated with CellEvent^™^ Senescence Green Probe (ThermoFisher Scientific, Waltham, MA) according to the manufacturer’s instruction. Data were acquired using a LSR II flow cytometer.

### Transcriptome analysis using RNA sequencing

For marrow HSPC RNA sequencing (RNA-seq), wild-type (CD45.1) and JAK2V617F mutant (CD45.2) marrow Lin^-^cKit^+^ HSPCs were isolated by magnetic bead isolation (with 90-95% purity) (Miltenyi Biotec, San Diego, CA). Briefly, the Lineage Cell Depletion Kit (Miltenyi Biotec) was used to deplete mature hematopoietic cells. The lineage negative cells were collected and then positively selected for CD117^+^ (cKit^+^) cells using CD117 microbead (Miltenyi Biotec) to yield Lin^−^cKit^+^ HSPCs. Messenger RNA samples of wild-type Lin-cKit+ HSPCs transplanted alone (one pooled sample from 3 mice), JAK2V617F mutant HSPCs transplanted together with wild-type cells (one pooled sample from 3 mice), and JAK2V617F mutant HSPCs transplanted alone (one pooled sample from 2 mice) were assessed by RNA-seq as we previously described^20^. Gene Ontology (GO) (http://www.geneontology.org/) enrichment analysis of differentially expressed genes was implemented by the ClusterProfiler R package. GO terms with corrected p values < 0.05 were considered significantly enriched.

### Statistical Analysis

Statistical analyses were performed using Student’s unpaired, 2-tailed *t* tests using Excel software (Microsoft). A *p* value of less than 0.05 was considered significant. Data are presented as mean ± standard error of the mean (SEM).

## RESULTS

### MPN disease relapse in a JAK2V617F-bearing vascular niche following lethal irradiation and marrow transplantation

Previously, we reported that a JAK2V617F-bearing vascular niche not only promotes the expansion of JAK2V617F mutant cells in preference to wild-type cells^21,22^ but also protects mutant HSPCs from irradiation or cytotoxic chemotherapy (e.g. busulfan)^17,23^, the two treatments commonly used in the conditioning regimen for stem cell transplantation in MPN patients. To investigate the underlying mechanisms for MPN disease relapse, we transplanted wild-type CD45.1 marrow cells directly into lethally irradiated (950cGy) Tie2FF1 mice or age-matched littermate control mice (CD45.2) (Figure 1A). During a ∼6-7mo follow up, while all wild-type control recipients displayed full donor engraftment, ∼60% Tie2FF1 recipient mice displayed recovery of the JAK2V617F mutant hematopoiesis (mixed donor/recipient chimerism) 10 weeks after transplantation, results consistent with our previous report^17^ (Figure 1B). In contrast to the Tie2FF1 recipient mice with full donor engraftment, the mixed chimeric mice developed neutrophilia and thrombocytosis (Figure 1C), similar to what has been observed in the primary Tie2FF1 mice^22,24^. Flow cytometry analysis revealed that marrow Lin^-^cKit^+^Sca1^+^CD150^+^CD48^-^ HSCs were significantly expanded in the Tie2FF1 recipient mice with mixed chimerism (i.e. with disease relapse) compared to Tie2FF1 recipient mice with full donor engraftment (i.e. with disease remission) (Figure 1D). Using CD45.1 as a marker for wild-type donor and CD45.2 for JAK2V617F mutant recipient cells, we found that the wild-type HSCs were severely suppressed and the JAK2V617F mutant HSCs were significantly expanded (0.001% vs. 0.128%, *P*=0.003) in the relapsed Tie2FF1 recipient mice; in contrast, there was no significant difference between the wild-type and mutant HSC numbers in the remission mice (0.023% vs. 0.012%, *P*=0.411) (Figure 1E).

**Figure 1.**
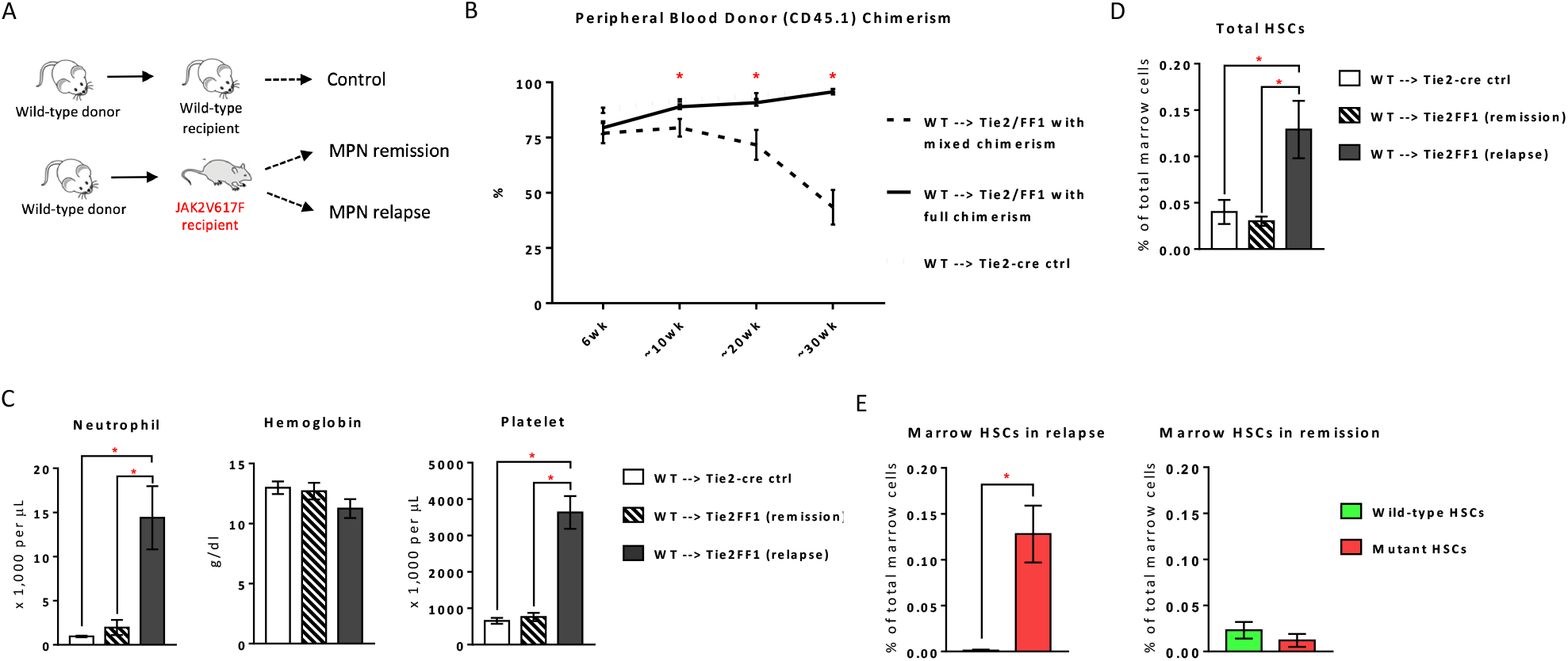
MPN disease relapse in a JAK2V617F-bearing vascular niche following lethal irradiation and marrow transplantation. (**A**) A murine model of MPN disease relapse established by marrow transplantations. (**B**) Peripheral blood CD45.1 chimerism following transplantation of wild-type (WT) CD45.1 marrow cells into lethally irradiated Tie2FF1 mice or Tie2-cre control mice (CD45.2) (n=8-12 mice in each group). (**C**) Tie2FF1 recipients with mixed chimerism developed both neutrophilia and thrombocytosis (n=5-8 mice in each group). (**D**) Total marrow Lin^-^cKit^+^Sca1^+^CD150^+^CD48^-^ HSCs (n=4-9 mice in each group). (**E**) Wild-type and JAK2V617F mutant HSCs in relapsed Tie2FF1 recipient mice (left) and remission Tie2FF1 recipient mice (right) (n=4-6 mice in each group). * *P* < 0.05

### The competition between wild-type and JAK2V617F mutant HSPCs during MPN disease relapse after marrow transplantation

Cell competition is an evolutionarily conserved mechanism involved in development, tissue homeostasis, and stem cell maintenance. It is similar to natural selection between species, in that “fitter” cells win out over their “less-fit” neighbors^25-27^. In humans, cells carrying oncogenic mutations that are phenotypically silent for many years are not uncommon. It has been suggested that cell competition can protect against the emergence of cancer^28,29^. Recently, we reported that exposure to wild type cells alters both the gene expression profile and cellular function of JAK2V617F mutant HSPCs and the presence of wild-type cells can prevent the expansion of co-existing JAK2V617F mutant cells^20^. We hypothesize that competition between the wild-type donor cells and JAK2V617F mutant recipient cells dictates the outcome of disease relapse versus remission after stem cell transplantation.

To test this hypothesis, we compared wild-type and JAK2V617F mutant HSPC functions between the relapsed Tie2FF1 recipient mice and those remained in remission. First, we measured wild-type and JAK2V617F mutant Lin^-^cKit^+^Sca1^+^ (LSK) cell proliferation *in vivo* by BrdU labeling. We found that the JAK2V617F mutant LSK cells proliferated more rapidly than wild-type LSKs during MPN relapse (45% vs. 9%, *P*=0.022), while there was no significant difference in cell proliferation between the mutant and wild-type LSK cells during remission (62% vs. 57%, *P*=0.805) (Figure 2A).

**Figure 2.**
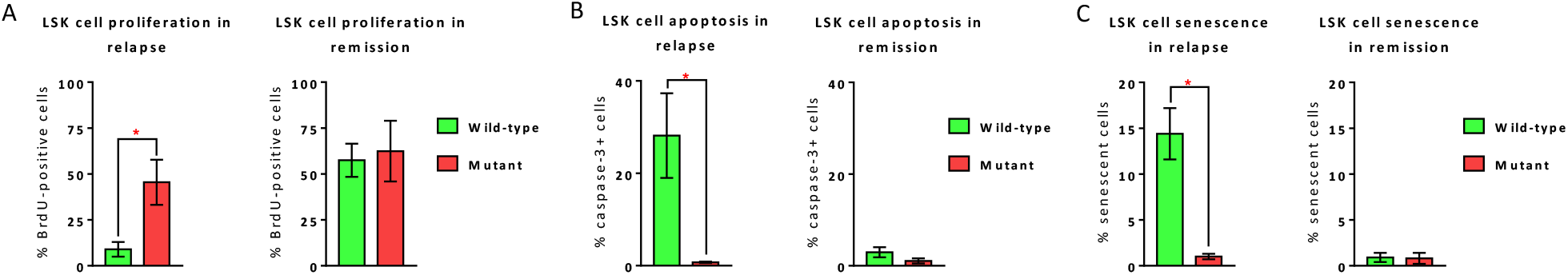
Comparison of wild-type and JAK2V617F mutant HSPC functions between relapse and remission. (**A**) Lin^-^cKit^+^Sca1^+^ (LSK) cell proliferation rate in relapse (left) and remission (right) Tie2FF1 recipient mice, measured by *in vivo* BrdU labeling (left: n=6 mice in each group; right: n=4 mice in each group). (**B**) Cellular apoptosis rate of wild-type and JAK2V617F mutant LSKs in relapse (left) and remission (right) Tie2FF1 recipient mice, measured by activated caspase-3 staining using flow cytometry analysis (left: n=4-6 mice in each group; right: n=3-5 mice in each group). (**C**) Cellular senescence rate of wild-type and JAK2V617F mutant LSKs in relapse (left) and remission (right) Tie2FF1 recipient mice, measured by SA-β-Gal activity using flow cytometry analysis (left: n=6 mice in each group; right: n=4-5 mice in each group). * *P*<0.05

We then measured LSK cell apoptosis by assessing activated caspase-3 staining. Wild-type LSK cells exhibited significantly higher level of apoptosis than mutant LSKs during relapse (28% vs. 0.7%, *P*=0.005); in contrast, there was no significant difference in the levels of cell apoptosis between wild-type and JAK2V617F mutant LSKs during remission (3% vs. 1%, *P*=0.286) (Figure 2B).

As the JAK-STAT signaling has been reported to induce cellular senescence^30-33^, we assessed cellular senescence by measuring senescence associated β-galactosidase (SA-β-Gal) activity using flow cytometry analysis. While wild-type LSKs demonstrated significantly higher senescence rates compared to JAK2V617F mutant LSKs during MPN relapse (14% vs. 1%, *P*<0.001), there was no difference in the cellular senescence rate between wild-type and mutant LSKs during remission (0.9 vs. 0.8%, *P*=0.941) (Figure 2C).

Taken together, there were dynamic alterations of the wild-type and JAK2V617F mutant HSPC functions in the Tie2FF1 recipient mice after the stem cell transplantation: in recipient mice who achieved disease remission, there was no significant difference in cell proliferation, apoptosis, or senescence between wild-type and JAK2V617F mutant HSPCs; in contrast, in recipient mice who relapsed after the transplantation, wild-type HSPC functions were significantly impaired (i.e., decreased proliferation, increased apoptosis, and increased senescence) which could alter the competition between co-existing wild-type and mutant cells and lead to the outgrowth of the JAK2V617F mutant HSPCs and disease relapse. These findings suggest that wild-type cells can prevent the relapse of MPN neoplastic hematopoiesis after stem cell transplantation.

### Immune regulation associated with wild-type and JAK2V617F mutant cell competition

To understand how co-existing wild-type cells prevent the expansion of JAK2V617F mutant HSPCs, we established a murine model of wild-type and JAK2V617F mutant cell competition^20^. In this model, when 100% JAK2V617F mutant marrow cells are transplanted alone into lethally irradiated wild-type recipients, the recipient mice develope a MPN phenotype with leukocytosis and thrombocytosis ∼4wks after transplantation; in contrast, when a 50-50 mix of mutant and wild-type marrow cells are transplanted together into lethally irradiated wild-type recipient mice, the JAK2V617F mutant donor cells engraft to a similar level as the wild-type donor cells, with the recipient mice displaying normal blood counts during more than 4-months of follow up^20^ (Figure 3A). To determine the underlying molecular pathways that could be differentially activated by wild-type and mutant cell competition, we performed gene expression profiling of JAK2V617F mutant Lin^-^cKit^+^ HSPCs with and without wild-type cell competition using RNA sequencing (RNA-seq). Gene ontology terms humoral immune response, leukocyte/B cell/T cell/complement activation, and immune response-activated signaling transduction were highly enriched in mutant HSPCs with cell competition compared to mutant HSPCs without competition (Figure 3B). These results prompted us to examine various immune cell types using flow cytometry analysis. We found that: (1) compared to wild-type HSPCs, JAK2V617F mutant HSPCs generated significantly more T cells and less B cells in the spleen, and more myeloid-derived suppressor cells (MDSCs)^34^ in the marrow; and (2) there was no difference in T, B, or MDSC numbers between recipients of wild-type HSPCs and recipients of mixed wild-type and JAK2V617F mutant HSPCs (Figure 3C). These data suggest that the JAK2V617F mutant stem cells have altered differentiation potential towards various immune cell types, and this abnormal differentiation can be “corrected” by the co-existing wild-type cells.

**Figure 3.**
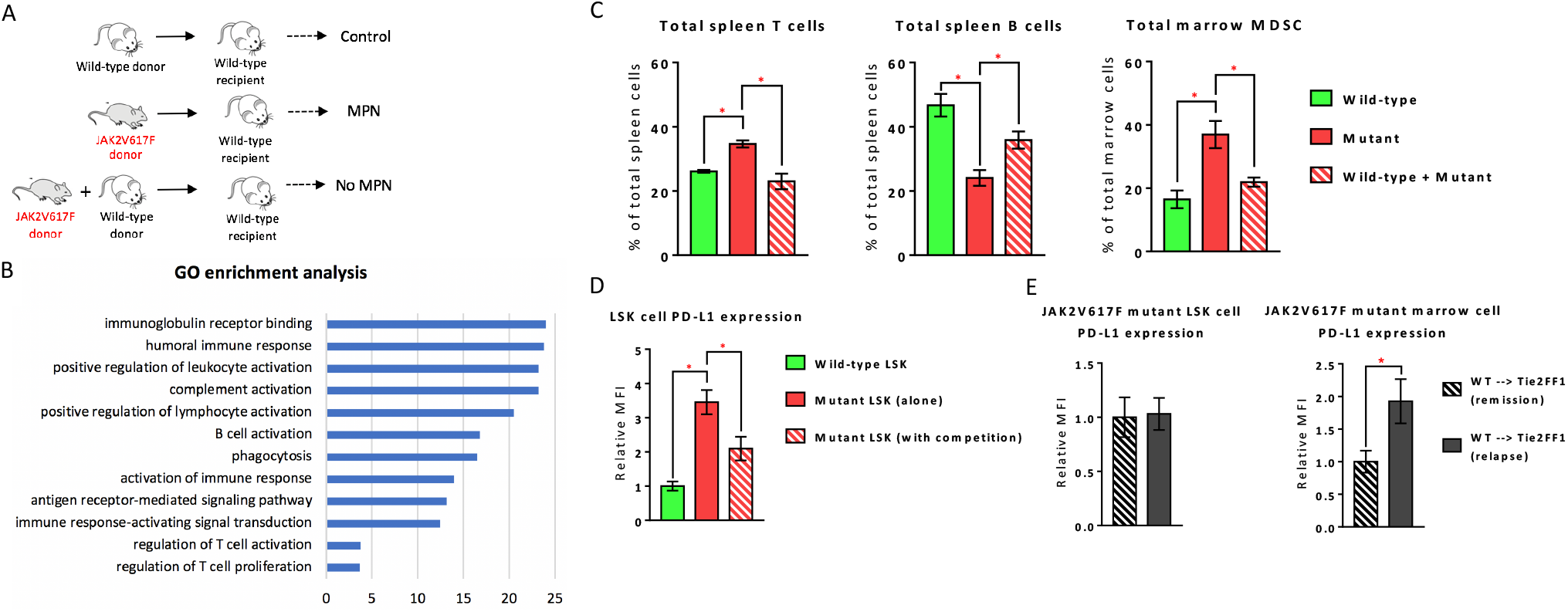
Immune regulation associated with wild-type and JAK2V617F mutant cell competition. (**A**) A murine model of wild-type and JAK2V617F mutant cell competition established by marrow transplantations. (**B**) Differentially enriched Gene Ontology (GO) terms in mutant Lin^-^cKit^+^ HSPCs transplanted together with wild-type cells (pooled sample from 3 mice) compared to mutant HSPCs transplanted alone (pooled sample from 2 mice). *P* values are plotted as the negative of their logarithm. (**C**) Spleen T cells (CD3^+^CD4^+^ and CD3^+^CD8^+^) and B cells (CD3^-^B220^+^), and marrow MDSCs (both CD11b^+^Ly6C^high^Ly6G^-^ M-MDSCs and CD11b^+^Ly6C^low^Ly6G^+^ PMN-MDSCs) in wild-type recipient of wild-type donor (“wild-type”), JAK2V617F mutant donor (‘mutant”), or both wild-type and mutant donors (“wild-type + mutant”) (spleen T cells: n=3 mice in each group; spleen B cells: n=3-6 mice in each group; marrow MDSCs: n=3-6 mice in each group). (**D**) Quantitative measurement of mean fluorescence intensity for PD-L1 staining on wild-type LSK cells (n=7 mice), JAK2V617F mutant LSK cells (n=4 mice), and JAK2V617F mutant LSK cells with co-existing wild-type cell competition (n=9 mice). (**E**) Quantitative measurement of mean fluorescence intensity for PD-L1 staining on JAK2V617F mutant LSK cells (left) and unfractionated marrow cells (right) during disease relapse and remission (n=5-6 mice in each group). * *P*<0.05

Program death ligand 1 (PD-L1) is expressed by many tumor cells. It binds to program death receptor-1 (PD-1) expressed on activated T cells and results in T cell exhaustion, thereby blunting the anti-tumor immune response and promoting tumor growth^35^. Recent studies demonstrated that the JAK2V617F mutation not only induces specific T cell response as a cancer neoantigen^36^ but also increases PD-L1 expression which can be used by the neoplastic cells to evade the antitumor immune response^37,38^. We did not detect any difference in PD-L1 gene expression between the wild-type and JAK2V617F mutant HSPCs using RNA-seq. Therefore, we measured cell surface PD-L1 protein expression by flow cytometry. We found that: (1) PD-L1 expression was significantly upregulated on JAK2V617F mutant LSK cells (3.5-fold, *P*<0.001) compared to wild-type cells; (2) PD-L1 expression on mutant LSK cells with co-existing wild-type cell competition was significantly decreased compared to mutant LSKs without cell competition (0.6-fold, *P*=0.009) (Figure 3D). These data suggest that competition between wild-type and JAK2V617F mutant cells triggered a potential immune regulation with dynamic changes in PD-L1 expression on the mutant HSPCs caused by their interactions with neighboring wild-type cells. We then tested whether such immune regulation also played a role in MPN disease relapse after stem cell transplantation (Figure 1A). We found that, although there was no difference in PD-L1 expression on JAK2V617F mutant LSK cells between relapse and remission, PD-L1 expression was significantly downregulated on JAK2V617F mutant unfractionated marrow cells in remission (where there were equal numbers of wild-type and mutant HSCs) compared to those in relapse (where wild-type HSCs were severely suppressed and the mutant HSCs were significantly expanded) (0.5-fold, *P*=0.012) (Figure 3E). Therefore, the presence of wild-type cells induced PD-L1 downregulation during both MPN clonal expansion and disease relapse.

Taken together, these results indicate that the wild-type cells may prevent the expansion of co-existing JAK2V617F mutant cells through modulating the immune abnormality induced by the JAK2V617F mutation.

## DISCUSSION

MPN relapse occurs in up to 40% of patients following stem cell transplantation (especially after reduced intensity conditioning)^5,7-9,39^. Many patients with MPNs or other hematologic malignancies develop mixed donor/recipient chimerism after allogenic stem cell transplantation. Some patients with such mixed chimerism can survive for a long time without disease relapse and there appears to be no difference in overall survival between patients with mixed donor/recipient chimerism and patients with full donor chimerism^40-45^. The implications of such mixed chimerism in MPN disease relapse is not well understood.

Cell competition is an evolutionarily conserved mechanism in which “fitter” cells out-compete their “less-fit” neighbors. It is seen only when there is a mixture of genetically different cells, a phenomenon known as mosaicism. Human tissues exhibit high levels of mosaicism with many precancerous mutations, yet such tissues rarely develop frank tumors^46,47^. Recently, we reported that the presence of wild-type cells alters both the gene expression profile and cellular function of JAK2V617F mutant HSPCs and inhibits the expansion of co-existing JAK2V617F mutant cells^20^. Here, using a MPN disease relapse murine model^17^ (Figure 1), we have been able to highlight the importance of cell competition between wild-type (donor) and mutant (recipient) cells during MPN disease relapse after stem cell transplantation. Specifically, we showed that wild-type HSPCs from the relapsed mice demonstrated impaired proliferation and increased apoptosis and senescence compared to wild-type HSPCs from the remission mice; in contrast, there was no significant difference in mutant HSPC functions (i.e. proliferation, apoptosis, senescence) between relapse and remission (Figure 2). These findings suggest that deterioration of wild-type cell function is important for MPN disease relapse. One limitation of this study is that we only examined HSPC function at 7-8 months after transplantation. Whether impairment of wild-type HSPC function occurs before disease relapse or is a consequence of MPN disease relapse is not clear, although these two mechanisms are not necessarily exclusive.

Patients with MPNs are characterized by a significant dysregulation of the immune system^36,37,48,49^. A number of recent studies have characterized the immune microenvironment of human solid tumors, where malignant transformation coincides with extensive remodeling of the immune microenvironment^50-52^. Immune normalization of the tumor microenvironment has contributed to successful anti-tumor efficacy of immunotherapies targeting the PD-1/PD-L1 pathway^53^. We demonstrated here that the presence of co-existing wild-type cells can correct the abnormal immune cell composition of the hematopoietic microenvironment and downregulate the PD-L1 expression of the JAK2V617F mutant HSPCs (Figure 3). These findings suggest that wild-type cells can restore the immune dysregulation induced by the JAK2V617F oncogene. Further investigation of tumor-specific immune response in the hematopoietic microenvironment is required to understand how the immune system contribute to the regulation of cell competition between wild-type and JAK2V617F mutant cells.

## CONCLUSION

In summary, although the molecular mechanisms responsible for MPN disease relapse after stem cell transplantation remain unclear, our study provides the important observations and mechanistic insights that co-existing wild-type cell competition can prevent MPN disease relapse after stem cell transplantation, the only curative treatment for patients with these diseases. Results from our previous work^20^ and current study provide the rational to further investigate whether wild-type cells could be used as a therapeutic approach to control mutant clonal expansion in MPNs. It also calls into question the best therapeutic approach for patients with mixed donor/recipient chimerism after transplantation, where timing might be more important than speed.

## DECLARATIONS

### Ethics approval and consent to participate

Animal experiments were performed in accordance with the guidelines provided by the Institutional Animal Care and Use Committee.

### Consent for publication

Not applicable.

### Availability of data and materials

Upon request to the corresponding author

### Author contributions

H. Zhang performed various *in vitro* and *in vivo* experiments of the project; M.C. assisted various *in vivo* experiments; L.Z. provided scientific consultation on the immune phenotype associated with wild-type and JAK2V617F mutant cell competition; H. Zhan conceived the projects, analyzed the data, interpreted the results, and wrote the manuscript.

